# Discovery of signatures of fatal neonatal illness in vital signs using highly comparative time-series analysis

**DOI:** 10.1101/2021.03.26.437138

**Authors:** Justin C Niestroy, J Randall Moorman, Maxwell A Levinson, Sadnan Al Manir, Timothy W Clark, Karen D Fairchild, Douglas E Lake

## Abstract

**Objective:** To seek new signatures of illness in heart rate and oxygen saturation vital signs from Neonatal Intensive Care Unit (NICU), we implemented highly comparative time-series analysis to discover features of all-cause mortality in the next 7 days.

**Design:** We collected 0.5Hz heart rate and oxygen saturation vital signs of infants in the University of Virginia NICU from 2009 to 2019. We applied 4988 algorithmic operations from 11 mathematical families to random daily ten-minute segments. We clustered the results and selected a representative from each, and examined multivariable logistic regression models.

**Setting:** Neonatal ICU

**Patients:** 5957 NICU infants; 205 died.

**Measurements and main results:** 3555 operations were usable; 20 cluster medoids held more than 81% of the information. A multivariable model had AUC 0.83. Five algorithms outperformed others: moving threshold, successive increases, surprise, and random walk. We computed provenance of the computations and constructed a software library with links to the data.

**Conclusions:** Highly comparative time-series analysis revealed new vital sign measures to identify NICU patients at the highest risk of death in the next week.

## Introduction

Continuously monitored vital signs of patients in intensive care units hold untapped information about risk for adverse events and outcomes ^1^. For example, the display of a score based on analysis of abnormal heart rate characteristics was shown by our group to reduce sepsis-associated mortality by 40% in preterm infants in the Neonatal Intensive Care Unit (NICU) ^2–5^. That approach was tailored to detect specific phenomena that we observed in the heart rate data, reduced variability and transient decelerations, in the days prior to sepsis diagnosis^2–5^, and we used algorithms optimized for the task, including sample asymmetry ^6^ and sample entropy ^7–9^.

Here, we asked a more general question - what if we did not know all the characteristics we wish the algorithms to detect? That is, if we used a very large number of algorithms designed for general use in time-series, would we discover some that were more effective than our tailored design? This approach has been described by Fulcher and coworkers, who called it *highly comparative time-series analysis* ^10–12^. The fundamental idea is to extract features from many time series, using many algorithms, most operating with many sets of parameter values. We then apply this ensemble to a data set to determine which algorithms perform best for predicting a specific outcome. Clustering of algorithms can, eventually, simplify this approach for clinical applications. ^13,14^

As an example of our approach, the familiar sample entropy algorithm ^7,8^ requires two parameters in order to operate, an embedding dimension *m* and a tolerance window *r*. A highly comparative time-series analysis entails many operations of the sample entropy algorithm that vary *m* and *r*. The result of each operation is treated as a potential predictor. Since the results are expected to be highly correlated, we can represent the family of sample entropy results as a cluster, and choose an operation of sample entropy with a single optimal combination of *m* and *r* for use in multivariable statistical models.

Furthermore, rather than simply clustering methods we know to be in the same family (“sample entropy” etc.), but with differing parameters, we can expand clustering to include many families of methods, and their parameters, defining the clusters using an outcome similarity measure. The cluster that contains sample entropy might then also contain related measures detected by clustering, all of which can be represented by a single outcome measure or feature. In all, this is an efficient way to screen many time-series algorithms, to discover features that are predictive of an outcome, without domain knowledge of prior known specific characteristics such as reduced heart rate variability, that might be related to that outcome.

To test these ideas, we selected death in the next 7 days for infants in the NICU as the outcome of interest. This is a topic of clinical interest and usefulness - identification of infants at high risk, especially where risk appears to be rising quickly, can alert clinicians to the possibility of imminent clinical deterioration from illnesses such as sepsis or respiratory failure. The heart rate characteristics score noted above, which is targeted toward a specific time-series phenotype, has modest performance in this area ^15^. In this work, the question is whether an examination - and potentially a combination - of many time series feature extraction algorithms may improve on this targeted approach.

## Materials and Methods

### Study design

We collected all bedside monitor data from all patients in the Neonatal ICU (NICU) at the UVA Hospital since 2009 using the BedMaster Ex™ system (Excel Medical, Jupiter FL). heart rate derived from the bedside monitor electrocardiogram signal is sampled at 0.5 Hz. oxygen saturation is measured using Masimo SET® pulse oximetry technology (Masimo Corporation, Irvine CA) with a sampling rate of 0.5 Hz and averaging time of 8 seconds. For this analysis, we included all infants admitted from 2009 through 2019 who had heart rate and oxygen saturation data available for analysis. Clinical data were abstracted from a NICU database (NeoData, Isoprime Corporation, Lisle, IL). This work was approved by the UVA Institutional Review Board with waiver of consent.

### Patient population and causes of death

From January 2009 through December 2019, 6837 infants were admitted to the UVa NICU, with median gestational age (GA) 35 weeks. Of these, 5957 infants had heart rate and oxygen saturation data available for analysis, and 205 died and had heart rate and oxygen saturation data available within 7 days of death. The primary causes of death (in seven categories) and associated clinical variables are shown in **Table 1**. For 152 of the 205 infants that died, support was redirected due to critical illness and grim prognosis. Of these, 148 died within minutes to hours after removal from mechanical ventilation. The other 4 infants died 2 to 4 days after the ventilator was discontinued, during which time comfort measures were provided. The remaining 53 infants died while on mechanical ventilation. In all cases, full support was provided while the infants were on mechanical ventilation.

**Table 1:** Families of algorithms implemented in highly comparative time series analysis.

| Family | Description | Example(s) |
| --- | --- | --- |
| Distribution | Moments and other descriptive statistics | Mean, median, standard deviation |
| Correlation | Similarity of data points as a function of the time between them | Linear and nonlinear autocorrelation |
| Stationarity | Statistical properties do not change over time | Standard deviation of moments measured on different window lengths |
| Symbolic transforms | Convert ranges to letters and analyze their sequence | Frequency of successive increases |
| Entropy | Order and regularity | Sample entropy |
| Trend analysis | Fitting lines through data | Slope and intercept |
| Heart Rate Variability | Canonical analyses | Power spectral density ratios |
| Time Series Modelling | Fits time series model to data | Surprise |
| Wavelet | Properties of the time series wavelet spectrum | Wavelet decomposition of time series |
| Nonlinear Analysis | Nonlinear analysis methods | False nearest neighbors, Information dimension |
| Other | Extreme values | Moving threshold model |

### Software and computing environment

We prepared a library of software in Python3 consisting of our implementations of 111 published algorithms ^10^ described for use in medical and non-medical domains. Table 1 shows the families of algorithms employed, together with a description and examples of each.

We ran these routines and some additional special-purpose MATLAB algorithms in Docker containers designed to run in a horizontally scalable secure cluster environment under the OpenStack cloud operating system, using the FAIRSCAPE data lake environment. ^16^ We issued persistent identifiers for all software, datasets and analysis results using Archival Resource Keys (ARKs) ^17^, associated with computational provenance metadata ^18^ for reproducibility and reusability.

### Terminology: Features, algorithms, and operations

A *feature* of a vital signs time series is a pattern or phenotype that can be represented mathematically. For example, we speak of the features of heart rate time series before neonatal sepsis as abnormal heart rate characteristics of reduced variability and transient decelerations. *Algorithms* are the mathematical tools we use to quantify the features. For example, the standard deviation of the times between heartbeats quantifies the finding of reduced heart rate variability in illness. *Operations* further specify the details of algorithms. For example, the standard deviations of the times between heartbeats over the past 5 seconds or 5 minutes or 5 hours all quantify heart rate variability, but they will return different values and, possibly, be of different utility clinically. The goal of highly-comparative time series analysis is to seek new features by widely exploring the spaces of algorithms and operations.

### Mathematical analysis of vital signs

The raw vital signs data were stored as vectors of time stamps 2 seconds apart with the corresponding measurement of heart rate or oxygen saturation. We grouped the vital signs data into more than 18 million 10-minute non-overlapping windows, each with 300 measured values. In each group, we computed 81 time-series algorithms with varying parameters for a total of 2499 operations. The result was a matrix of results with more than 18 million rows and 2499 columns, as illustrated in the workflow diagram in **Figure 1**.

**Figure 1.**
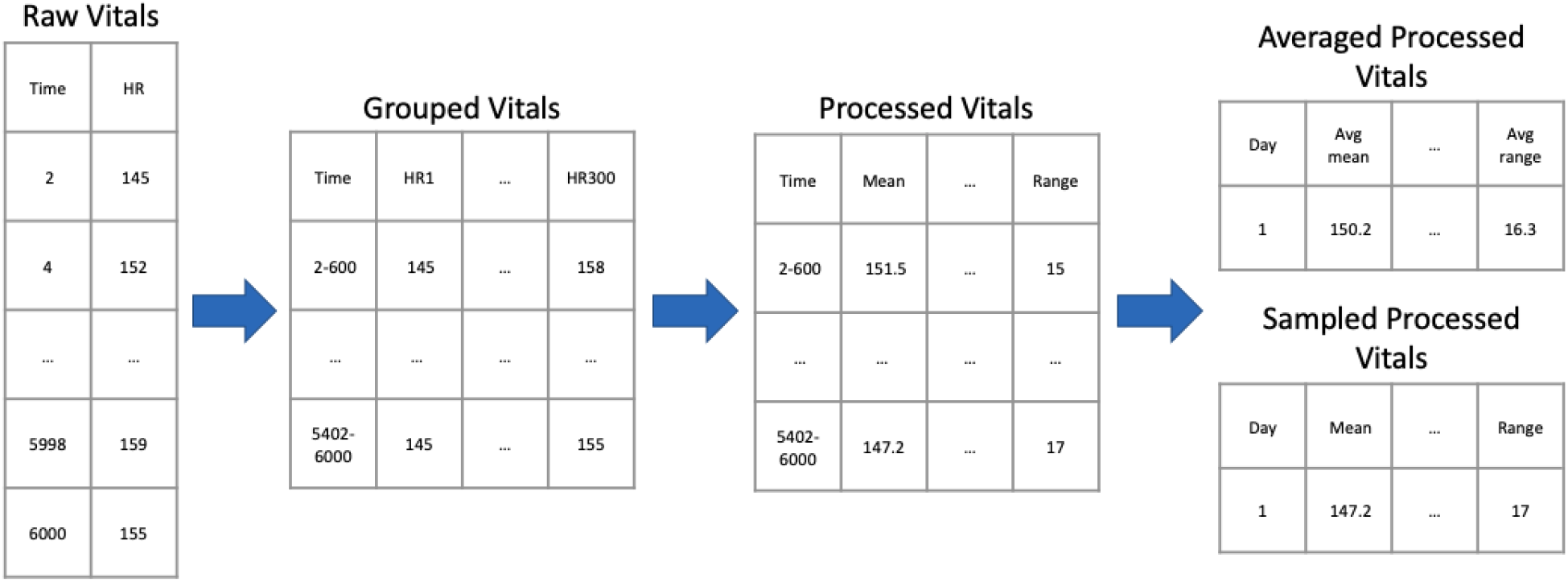
Data processing workflow for heart rate data. From left to right: Each row of the *Raw Vitals* table contains measured vital signs at 2 second intervals. These values are transposed in the *Group Vitals* table so that each row has a 600 second time range and up to 300 measured values. Algorithms operate on the *Group Vitals* table producing the *Processed Vitals* table of the same size - in the example, the first algorithm is the mean, and the last is the range. The *Averaged Processed Vitals* table holds the average of each result for a day; the *Sampled Processed Vitals* table holds the results for a randomly selected 600 second record.

We randomly sampled the *Processed Vitals* dataset taking one 10-minute record per day per patient. This step resulted in 130,000 days of samples, each containing the result of 4998 operations from the heart rate and oxygenation data. We removed single-valued results, those with imaginary numbers, and samples with missing values, and were left with 3555 of the 4998 viable candidate algorithmic operations. To adjust for the wide range in scales, we used an outlier-robust sigmoid transform ^10,19^ to convert operation ranges to the interval [0,1].

We clustered results to reduce dimensionality. We divided the 130,000 results of individual algorithms into ten equiprobable bins and calculated all possible distances using mutual information ^20 21^. We organized these results into a distance matrix and determined clusters with k-medoids using the *pam* function of the R *cluster* package ^22^. We represented each cluster by a single operation, as shown in **Figure 2**.

**Figure 2.**
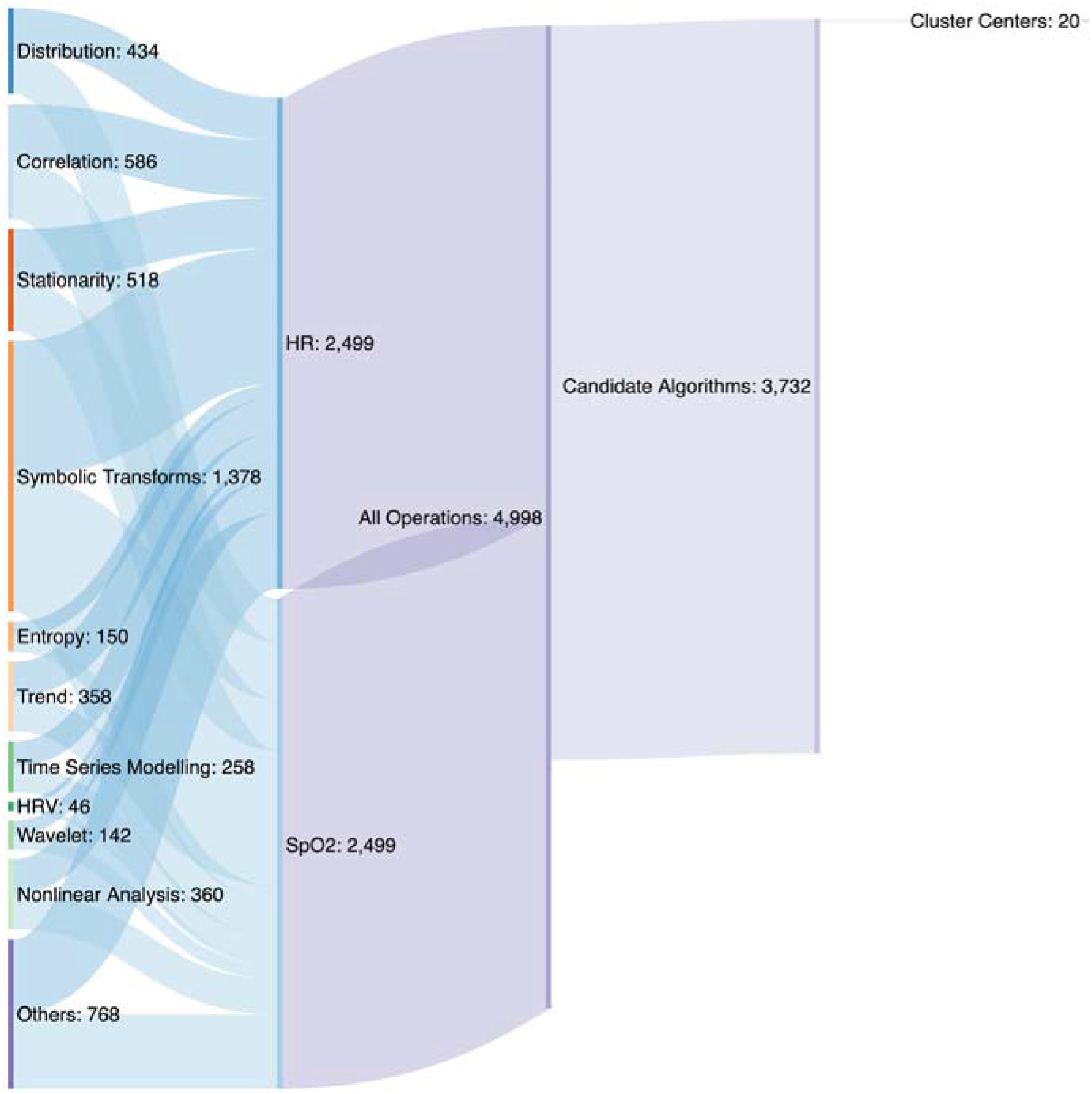
Sankey plot of the computational approach. Beginning with 3555 operations representing 11 families of algorithms, we ended with 20 clusters each represented by a single operation.

### Statistical analysis and modeling

The binary outcome of death within the next 7 days was used to evaluate algorithm performance. Since there were 205 deaths, we restricted the number of clusters to 20, and selected the top performers in each as candidate features for model selection. This follows recommendations to use no fewer than 10 events for each predictor variable. ^23^ Several feature selection strategies were used including lasso, greedy stepwise selection, AIC, and all-subset logistic regression. For simplicity and to be extra conservative to prevent over-fitting, we decided to concentrate on models with no more than 5 features. A stepwise backwards procedure was used that started with a full logistic regression model and sequentially removed features with largest p-value until there were 5 features. The performance of the model was calculated as the AUC using 10-fold cross-validation.

## Results

### Patient population

**Table 2** gives the demographics of the patient population and the causes of death.

**Table 2:** Demographics and diagnoses of the patient population.

| **Primary Cause of Death Categories** | ALL | Extreme prematurity first week deaths* | Brain disorders | Congenital cardiac disease | Multiple congenital anomalies** | Lung disease, pulmonary hypertension | Sepsis, Necrotizing Enterocolitis | OTHER*** |
| --- | --- | --- | --- | --- | --- | --- | --- | --- |
| n (%) | n=205 (100%) | n=26 (13%) | n=37 (18%) | n=23 (11%) | n=33 (16%) | n=38 (18%) | n=28 (14%) | n=20 (10%) |
| Gestational age (mean weeks) | 32.4 | 24.7 | 36.2 | 36.3 | 34.7 | 31.1 | 29.2 | 33.9 |
| Birth weight (mean kg) | 2.03 | 0.735 | 2.952 | 2.895 | 2.044 | 1.705 | 1.447 | 2.405 |
| Sex (%female) | 42% | 35% | 41% | 32% | 43% | 29% | 46% | 55% |
| Race (%Black) | 19% | 27% | 16% | 13% | 18% | 21% | 7% | 35% |
| Ethnicity (%Hispanic) | 4% | 4% | 3% | 4% | 12% | 0% | 4% | 0% |
| Age at death (mean) | 24.3 days | 2.1 days | 10.9 days | 10.6 days | 22.2 days | 54.6 days | 32.0 days | 29.0 days |
| Event Days | 871 | 64 | 149 | 87 | 140 | 182 | 153 | 96 |
| HR SPO2 Top 5+Demographic | 0.853 | 0.913 | 0.915 | 0.894 | 0.900 | 0.875 | 0.744 | 0.774 |
| HR SPO2 Top 5 | 0.828 | 0.911 | 0.823 | 0.899 | 0.837 | 0.860 | 0.759 | 0.779 |
| Demographic | 0.714 | 0.709 | 0.897 | 0.713 | 0.823 | 0.721 | 0.512 | 0.607 |
| *<29 weeks gestation with respiratory failure and/or severe intraventricular hemorrhage |  |  |  |  |  |  |  |  |
| **including genetic syndromes |  |  |  |  |  |  |  |  |
| ***including hydrops, metabolic disorders, renal or liver failure, and unknown |  |  |  |  |  |  |  |  |

### AUC for death prediction for each of the 3555 operations

In total, there were 871 daily ten-minute samples within a week of death for 205 infants, for a sample incidence rate of 0.67%. **Figure 3** shows the number of algorithms as a function of their univariate predictive performance for death in the next week, measured as AUC. The top performing algorithm, a symbolic logic count of successive increases in heart rate that is discussed further below, had an AUC of 0.799, substantially higher than that of the traditional algorithms like standard deviation of heart rate (0.749) and mean oxygen saturation (0.639).

**Figure 3.**
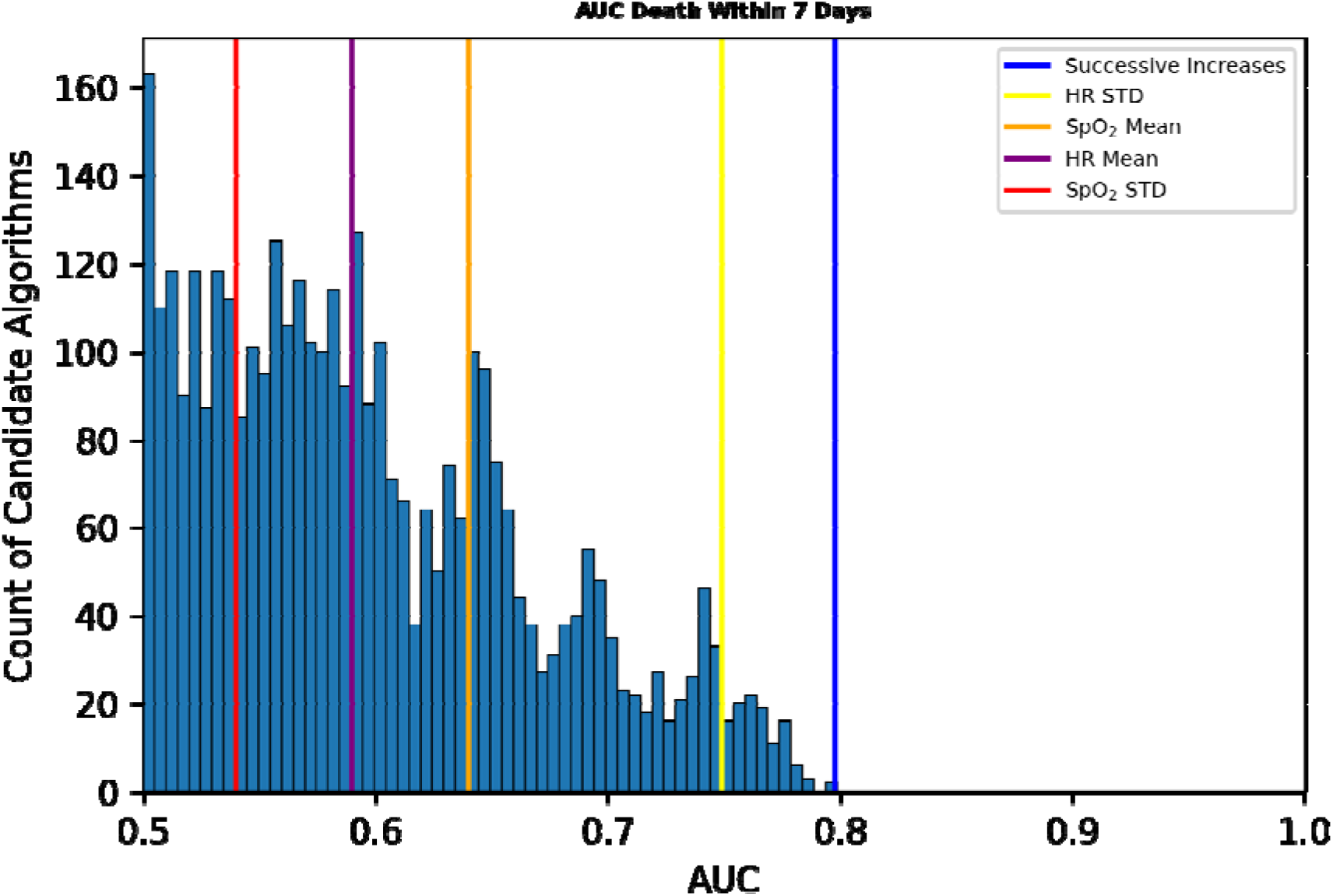
AUCs of 3555 operations for predicting death in the next 7 days. Colored vertical bars from left to right indicate the AUCs of the standard deviation of oxygen saturation, mean heart rate, mean oxygen saturation, standard deviation of heart rate, and a novel measure, successive increases of heart rate.

### Algorithmic results clustered, allowing data reduction

We sought correlations among the results. **Figure 4** shows two heat maps based on the absolute value of the correlation coefficients for 3555 algorithmic operations on the left and 20 identified by cluster medoids on the right.

**Figure 4.**
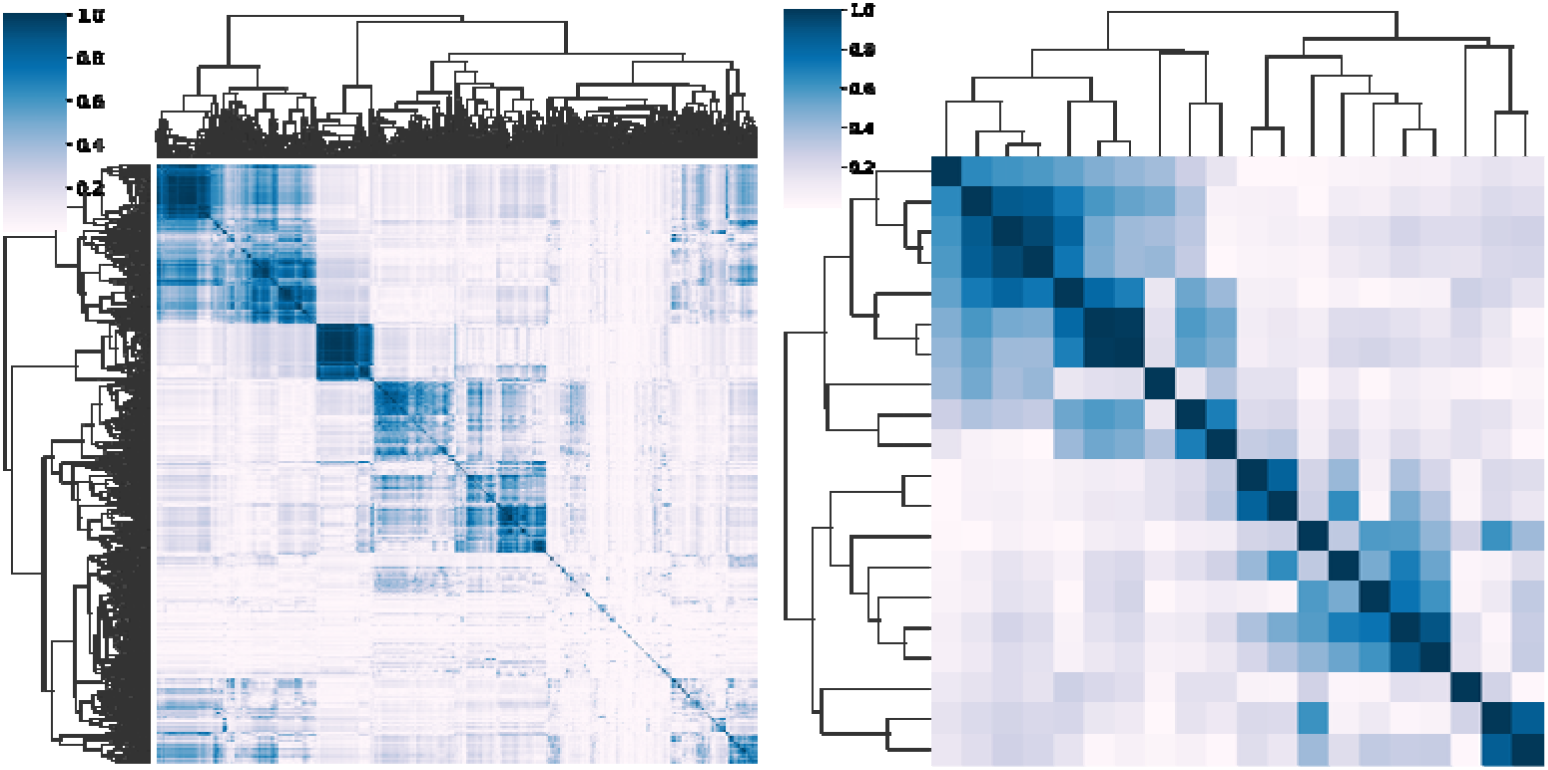
Heat maps of the absolute values of the correlation coefficients between results of operations. Left: correlations between all 3555 candidate algorithmic operations. Right: Correlations between 20 cluster medoids. The reduced feature set explains 81% of the variance in the full set.

These results justified an analysis of clusters of results, which we undertook by measuring mutual information among all the operational results. A representative result is shown in **Figure 5**. We sought a number of clusters that was large enough to explain most of the predictive performance of a multivariable statistical model for death but in keeping with the practice of having a reasonable number of predictors for 200 events. ^23,24^ We found that 20 clusters satisfied these conditions. The findings were robust in repeated experiments with different random sampling of one record per patient per day as well as daily averaged data.

**Figure 5.**
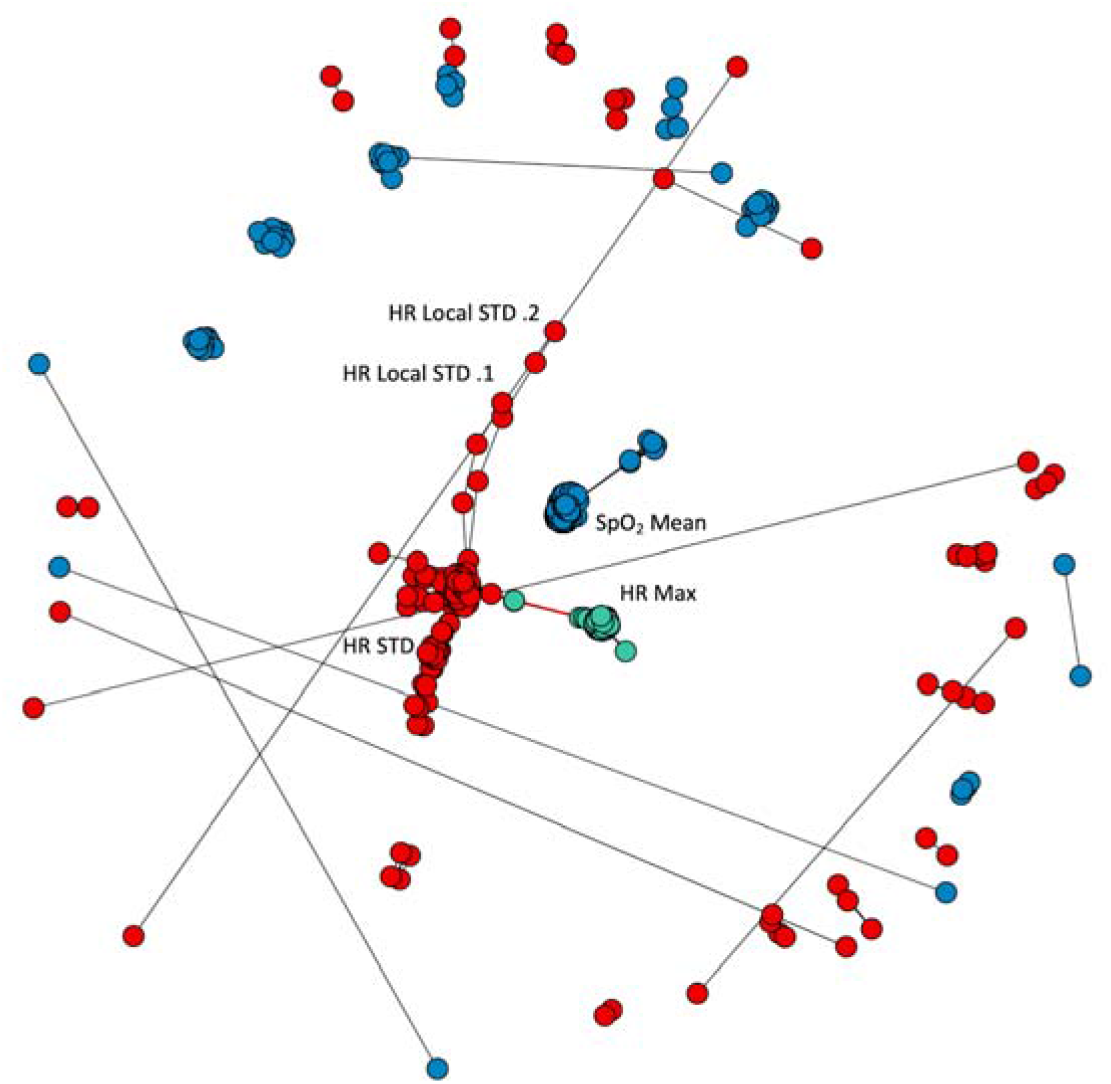
Three representative clusters of operations and their relations to each other. Each dot represents an individual operation. The colors - green, blue, and red - represent the three clusters; several individual operations are labeled. The green group represents measures of the maximum of the heart rate, the blue group reports on the mean oxygen saturation, and the red group reports on the standard deviation of the heart rate. The black lines indicate pairings between operations in the same cluster, whereas the very short red line indicates the small number of pairings between operations from different clusters. heart rate heart rate; STD standard deviation.

We examined the clusters for interpretability, and found that clusters of algorithmic operations reported on identifiable and interpretable time-series characteristics. For example, one cluster held the results of operations that report on the mean, another cluster held operations that report on the minimum, and so on. As expected, very near neighbors represented the results of operations that are closely related. To reduce the dimensionality of the data, each cluster can then be represented by a single operation and further analysis done on those 20.

### Multivariable statistical models using the new measures of vital signs to predict death

To understand the relationship of these 20 operations to the outcome of death, we first found the top univariate performers in each cluster as measured by AUC. Multivariable logistic regression models were develop using feature selection via backwards selection and model size limit of 5. Each patient was represented on each day by 20 results - 10 for each heart rate and each oxygen saturation time-series - calculated on a 10-minute record chosen at random once per day. In order to reduce the effects of possible outliers, results were winsorized by clipping low and high values of the results at 0.1% and 99.9% respectively. The purpose of the analysis was to identify time-series analytics that are new to the study of infant heart rates and oxygen saturations that were more effective predictors of death than commonly used measures. Since we picked the operations to be representative of the clusters, we ensured better interpretability of the results.

We made models with only heart rate features, only oxygen saturation features, and both heart rate and oxygen saturation features. The main results are summarized in **Table 2**. The multivariate HR-only model with 5 features had an AUC of 0.809 compared to the best heart rate univariate model, successive increases, which had an AUC of 0.799. The oxygen saturation only model with 5 features had an AUC of 0.765. The combined heart rate and oxygen saturation model with 5 features had an AUC of 0.828 – this was the best-performing 5-feature model. The AUC decreased slightly to 0.821 when using only the top 3 features. As a comparison, a combined heart rate and oxygen saturation model that was selected using AIC had 13 variables had a slightly improved AUC of 0.834.

**Table 2:** Model performances as a function of days until death. AUC for models using a single random daily 10-minute segment of heart rate (HR), oxygen saturation (SpO_2_), or both are shown at various intervals before death.

| Model Name | Candidate Features | Model Size | <= 7 days | <= 1 Day | 3 Days | 7 Days |
| --- | --- | --- | --- | --- | --- | --- |
| HR SPO2 Top 5+Demographic | 21 | 6 | 0.853 | 0.903 | 0.819 | 0.774 |
| HR SPO2 Top 5 | 20 | 5 | 0.828 | 0.892 | 0.794 | 0.747 |
| HR SPO2 Top 3 | 20 | 3 | 0.821 | 0.887 | 0.781 | 0.742 |
| HR SPO2 Cluster Centers | 20 | 5 | 0.819 | 0.873 | 0.790 | 0.763 |
| HR Top 5 | 10 | 5 | 0.809 | 0.864 | 0.770 | 0.750 |
| HR Successive Increases | 1 | 1 | 0.799 | 0.858 | 0.755 | 0.746 |
| HR Mean/SD SPO2 Mean/SD | 4 | 4 | 0.774 | 0.816 | 0.733 | 0.731 |
| SPO2 Top 5 | 10 | 5 | 0.765 | 0.818 | 0.752 | 0.694 |
| Demographic | 4 | 4 | 0.714 | 0.710 | 0.683 | 0.697 |

Another clinically relevant measure is accuracy of the model at a threshold equal to the event rate of 0.67%. In this case, the sensitivity is the same as the positive predictive value (PPV). For rare events it is informative to look at the ratio of this value to the event rate, or lift. The heart rate model had a sensitivity/PPV of 11.1% at this threshold for a lift of 16.5.

We tested how much discriminating capability was retained when we used the medoids of the 20 clusters versus the top performers in each. A combined heart rate and oxygen saturation model with 5 features from the 20 medoids had an AUC of 0.821, comparable to that obtained using the top performers. We conclude that each cluster can be effectively represented by a single operation.

Model performance can be improved by including baseline demographic information. A model consisting of birthweight, gestational age, sex and 5-minute Apgar core had an AUC of 0.714, and 5-minute Apgar score was the most predictive feature. Adding this demographic model to the combined heart rate and oxygen saturation model with 5 features increased the AUC from 0.828 to 0.853.

To better understand how these models performed leading up to the day of death, AUCs for each model were calculated daily from 7 days to 1 day prior to death, as shown in **Table 2**. For example, the AUC at 7 days was calculated excluding all data from the 6 days prior to death and only using the model output between 6 and 7 days. This excludes many deaths in first week of life and others that might be deemed to be expected. As expected, model performance is highest one day prior to death. The combined heart rate and oxygen saturation model with 5 features was one of best performers, with AUC of 0.892 the day before death and 0.747 a week prior to death.

### New measures of vital signs associated with NICU death

We found new measures of heart rate and oxygen saturation signals associated with NICU deaths. In the heart rate time-series, the top performing measures were fewer occurrences of *successive increases* (illustrated in **Figure 6**) and larger *surprise*. In the oxygen saturation time-series, the most predictive measure was a *moving threshold* calculation ^25^ that showed fewer extreme events was informative in both the heart rate and the oxygen saturation time-series. An algorithm fitting a random walk model to the oxygen saturation time-series detected declines in the oxygen saturation.

**Figure 6.**
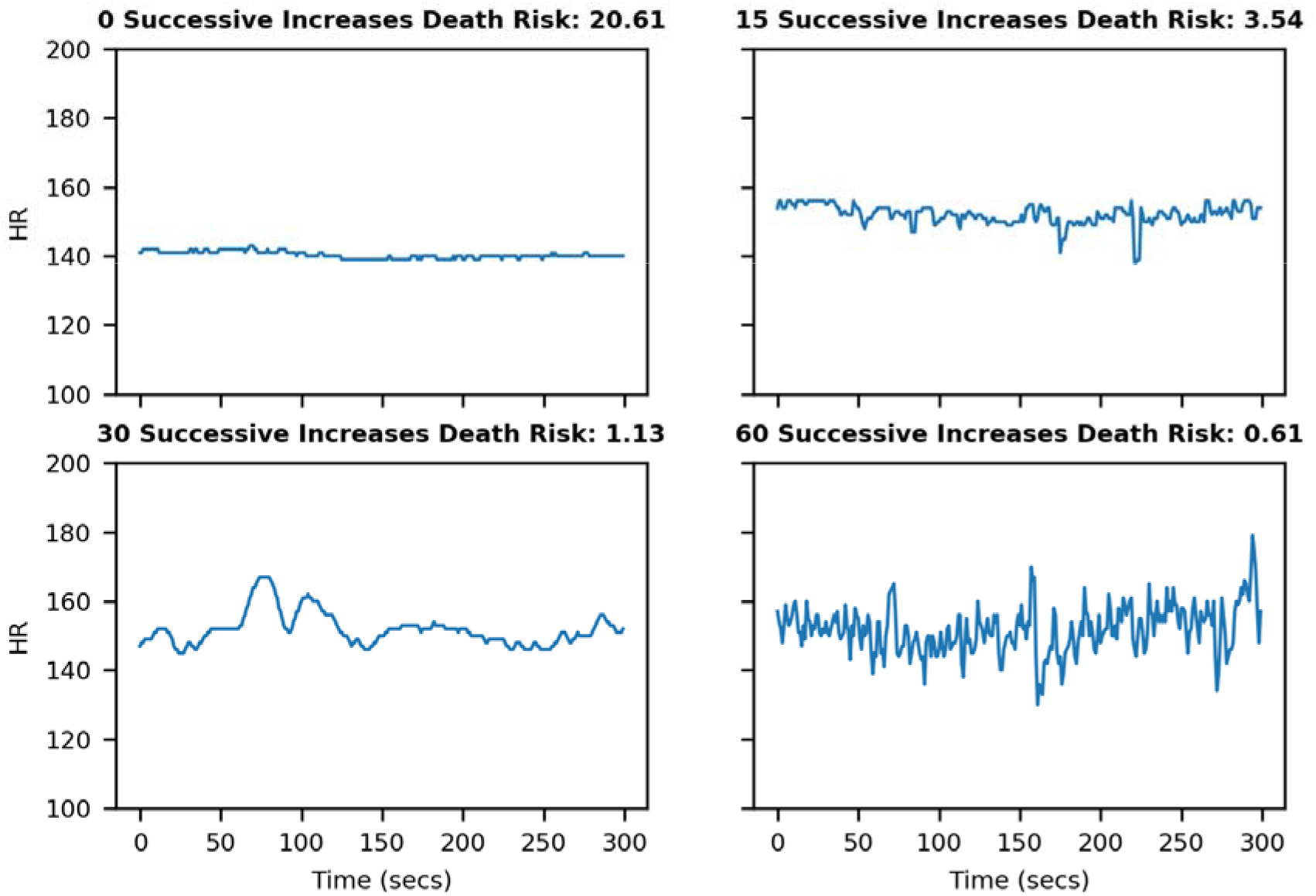
Lack of successive increases in heart rate predict increased risk of death. Four 10-minute heart rate records that correspond to increasing death risk with a decreasing number of *successive increases* in heart rate.

The most informative new predictor for death risk was a small number of successive increases in the heart rate, and **Figure 6** shows four records with increasing numbers of successive increases. The value that would be observed in a set of 300 random numbers, 75 (300/0.5^2^), is approached in the lower right panel. Qualitatively, the finding is that low heart rate variability is associated with higher risk of death. However, a more direct measure of variability, the standard deviation of the heart rate, was less predictive (AUC 0.799 for successive increases compared with 0.749 for heart rate STD).

## Discussion

Much progress has been made in the use of continuous time-series data from the bedside continuous cardiorespiratory monitors in the Neonatal ICU. ^26^ We tested the idea that we might improve the current art through a systematic study of our very large set of time series using an exhaustive number of analytical measures. We draw from a prismatic work describing the method of highly comparative time-series analysis, applying many time series algorithms to many time series examples of all kinds. ^10^ We applied the principles of highly comparative time series analysis to our domain, continuous cardiorespiratory monitoring in NICU patients. This work extends the study of highly comparative time-series analysis with its focus on clinical data sets, clinical events that are important to clinicians, and domain-specific knowledge of the physiologic origins of the data and how clinicians use it at the bedside.

In this example of highly comparative time-series analysis of a large clinical data set, we studied vital sign data from Neonatal ICU patients and discovered algorithms not previously reported in this domain that identified infants at higher risk of death. These algorithms report generally on the absence of heart rate variability and low oxygen saturation, features known to inform on poor health. Other major findings were that only 20 clusters of algorithms explained the great majority of the variance, in keeping with another study of highly comparative time-series analysis, ^14^ that downsampling of the data to single 10-minute records daily did not affect the overall results, and that the newly revealed algorithms outperformed standard measures of vital sign variability.

### A small number of clusters explain the variance of the results

This is not entirely unexpected, because many of the operations entail the same algorithm repeated with different arguments and parameters. For example, the sample entropy algorithm requires the choice of an embedding dimension *m* and a tolerance window *r*.^7–9^ In all, we performed 12 sample entropy operations with combinations of these variables. Thus, our clusters were to some extent explainable. For example, one held many operations that report on the center of the distributions, another held many reporting on the width of distributions, another held many entropy operations, and so on. These findings are important with regard to the interpretability of statistical models that use the results, avoiding the problem of black boxes. ^27^

### Downsampling of the data to single 10-minute records daily did not affect the overall results for algorithmic operations clustering

A downside to the massive exploration of algorithmic operations is the computing time. To begin our investigation, we accordingly massively reduced the data set to a single 10-minute record daily, <1% of the total. To test the fidelity of the results, we repeated the procedure on 3 other single 10-minute records, and on the daily average. The results were not significantly different in the nature of the clusters or their constituents, suggesting that a manageable subsample of the data can be used for exploratory purposes in the highly comparative time-series analysis, and the results verified afterward.

### Newly revealed algorithms outperformed canonical measures of vital sign variability

We discovered algorithms that out-performed the others but have not previously been applied to vital sign time series analysis. Importantly, they were interpretable in the light of domain knowledge about neonatal clinical cardiovascular pathophysiology.

*Successive increases* in heart rate is the result of a symbolic dynamics analysis, ^28^ and represent small accelerations. Individual vital sign measurements are replaced by symbols that reflect whether they have increased, decreased, or stayed the same compared to the preceding measurement. Our finding was that the number of consecutive increases in every-2-second heart rate was reduced in the time series of infants at higher risk of death, as illustrated in **Figure 6**. This finding is consonant with reduced heart rate variability, a known marker of abnormal clinical status. It is interesting to speculate why the absence of successive increases should improve upon ordinary measures of variability for prediction of death.

*Moving threshold* ^25^ was an approach developed in the field of extreme events in dynamical systems. An example is the human reaction to floods in rivers - when there are no floods, barriers are allowed to lapse. A flood, though, leads to new, higher barriers that could contain the recent event. The moving threshold model examines a time series for points that exceed an initial threshold, increases the threshold after a crossing event, and allows the threshold to decay over time. The parameters that are measured include the rate of events, the intervals between threshold crossings, and statistical measures of the distribution of the thresholds. Our finding was that the vital sign time series of infants at higher risk of death were characterized by lower moving threshold for heart rate (reflecting low heart rate variability) and lower moving threshold for oxygen saturation (illustrated in **Figure 7** as a gradual decline in SpO_2_).

**Figure 7.**
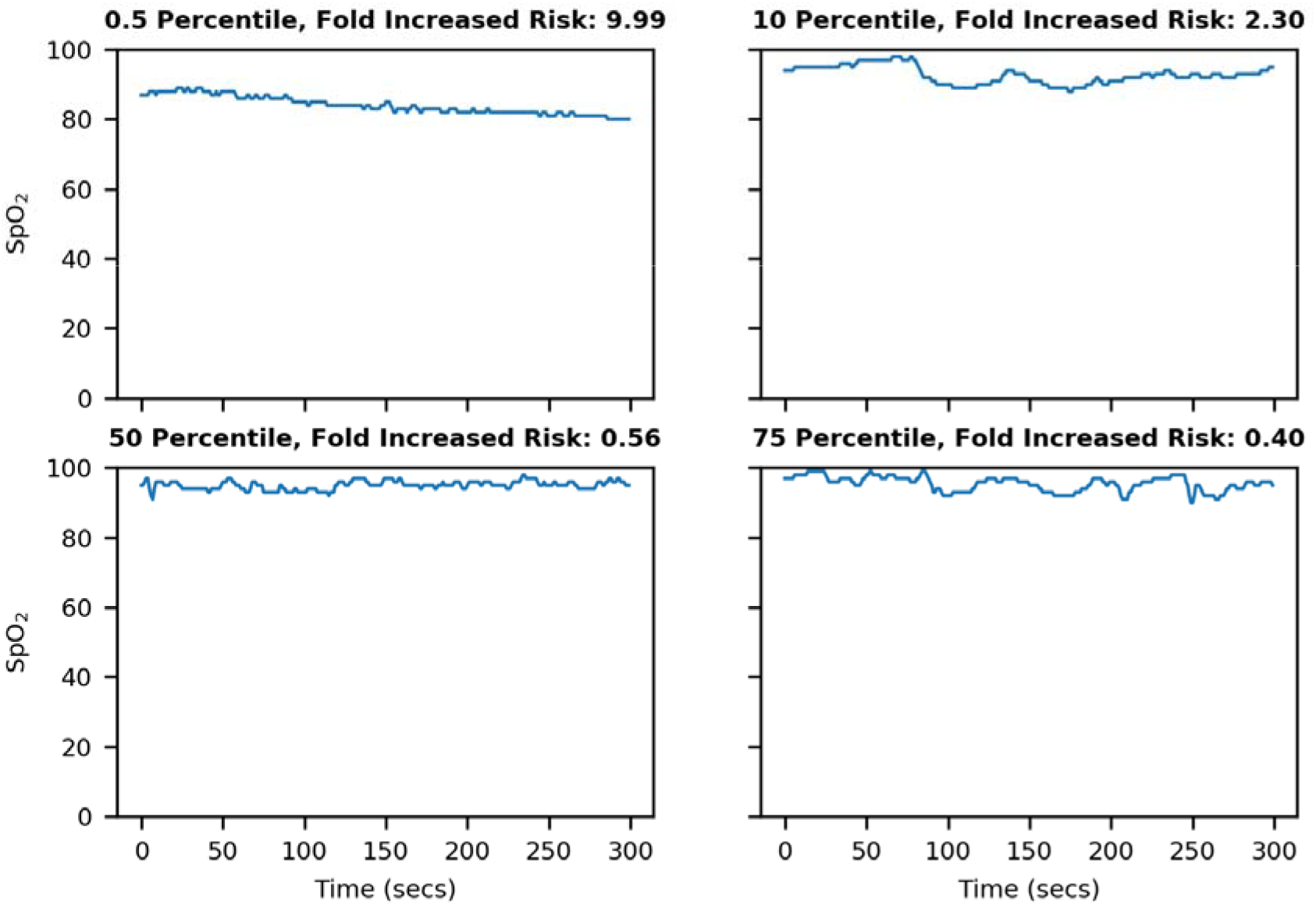
Lower moving threshold in oxygen saturation predicts increased risk of death. Four 10-minute heart rate records that correspond to increasing death risk with a decreasing percentile value for oxygen saturation moving threshold.

*Surprise* ^29^ calculates the distribution of points of a subsection of the time series. The surprise of the point following that subsection is measured by how likely the new point was given the calculated distribution, given as 1/*p*. The phenotype associated with mortality here, was a low value of surprise in the heart rate, consistent with reduced heart rate variability.

*Random walk model* measures the fit of the time-series data to a random walk. ^30^ The random walk starts at 0 and takes a step of size proportional to the distance from the previous point. The algorithm returns many statistics about the movement of the random walk and its relation to the original time series. The phenotype of high risk of death detected by this algorithm is a decline in the oxygen saturation.

### Relationship to prior work

In 2001, ^31^ we showed that heart rate characteristics of low variability and transient decelerations added information to clinical findings quantified by the SNAP (Score for Acute Neonatal Physiology) ^32^ and NTISS (Neonatal Therapeutic Intervention Scoring System) ^33^ in the early detection of sepsis. A heart rate characteristics index predicted sepsis and all-cause mortality in preterm NICU patients ^2,15,31,34^. We found that the AUC for the heart rate characteristics index developed at the University of Virginia and tested at Wake Forest University was 0.73. Subsequently we broadened the analyses to include conventional measures of heart rate andoxygen saturation in the first week after birth which we showed predict mortality among preterm NICU patients better than the validated and commonly accepted SNAPPE-II score (Score for Neonatal Acute Physiology-Perinatal Extension). ^35–37^ and combined heart rate and SpO_2_. ^38^ In 2010, ^39^ Saria and coworkers showed that short- and long-term variability of heart rate and oxygen saturation in the first 3 hours of life were useful in classifying premature infants at risk for high-morbidity courses. In the current work we showed that running an enormous number of operations on a single daily random 10 minute window of heart rate and oxygen saturation data uncovered new measures that predict mortality better than our prior models.

### Clinical implications

How might these findings lead to future improvements in neonatal care? We point to the increasing use of Artificial Intelligence and Machine Learning using Big Data to provide predictive analytics monitoring for early detection of subacute potentially catastrophic illnesses. While the data sources remain the same – continuous cardiorespiratory monitoring, lab tests, and vital signs measurements – the analytical methods are growing in number. An unresolved question has been whether the identification of signatures of illness by domain experts can be replaced by exhaustive computer analysis of large data sets. ^40^ These new findings point clearly to a role for highly-comparative time series analysis to detect previously unthought-of ways to characterized the pathophysiological dynamics of neonatal illness. Future work will test these new candidate algorithms against existing ones of heart rate characteristics analysis, ^41^ cross-correlation of heart rate and oxygen saturation, ^42,43^ heart rate variability, ^44^ and others.

### Limitations

We did not analyze pre-term infants separately from term infants in this work, though we know that heart rate and SpO2 time series characteristics depend on both gestational age and post-conceptual age. For example, the variabilities of heart rate and SpO2 rise with day of age, ^45,46^ and it is possible that highly-comparative time series analysis of pre-term infants might return different results from term infants. As it stands, there were many more term infants than pre-term, but the latter represented more of the time series data.

External validation will be important because our findings of patterns in vital signs measurements prior to neonatal death might reflect care practices at our hospital. We note, though, the similarity of vital signs measurements at our hospital to those at two others, ^45^ a finding that is reassuring with regard to the general nature of these results.

We found that only 3555 of the 4998 algorithms consistently returned non-null results, and we note that other data sets from other sources might fare differently. This was expected as many of the algorithms were from different domains and may not work on all signals.

We used only logistic regression to test the association of the algorithmic operations with clinical outcome, and other machine learning and deep learning methods might have had different results. We note, though, recent works that point to a similarity of results of logistic regression compared to other methods including recurrent neural networks. ^47,48^

## Conclusion

Highly comparative time-series analysis of clinical data reduced thousands of algorithms to an interpretable set of 20 that represent the character of neonatal vital signs dynamics.

This framework will be useful for future work analyzing bedside monitor data for signatures associated with various imminent or future adverse events and outcomes. The terabytes of vital sign data passing over monitors just at a single NICU such as ours, together with electronic medical record data on clinical and laboratory variables, hold valuable insights into actionable outcomes. Developing platforms and systems for sharing data with other investigators so that algorithms can be tested in large and diverse populations is another worthy goal. Harnessing these data could lead to preemptive strategies that improve patient outcomes.

## Data and Software Availability

Anonymized data that support the findings of this study, with the evidence graph for the clustering, are openly available in the University of Virginia’s LibraData archive at https://doi.org/10.18130/V3/VJXODP. Python code used to process this data is archived in Zenodo at https://doi.org/10.5281/zenodo.4321332. This version and any future versions are also available in Github at https://github.com/fairscape/hctsa-py. Our code is licensed under terms of the MIT license (https://opensource.org/licenses/MIT), and is a reimplementation in Python of most of Ben Fulcher’s original MATLAB code, available here: https://github.com/benfulcher/hctsa. Software for clustering analysis and cross-implementation testing, together with the test data, may be found here: https://doi:10.5281/zenodo.4627625.

